# Modeling the effect of spatial structure on solid tumor evolution and ctDNA composition

**DOI:** 10.1101/2023.11.10.566658

**Authors:** Thomas Rachman, David Bartlett, William Laframboise, Patrick Wagner, Russell Schwartz, Oana Carja

**Affiliations:** Computational Biology Department, School of Computer Science, Carnegie Mellon University, Pittsburgh, PA, USA; Joint Carnegie Mellon University-University of Pittsburgh Ph.D. Program in Computational Biology; Allegheny Cancer Institute, Allegheny Health Network, Pittsburgh PA

**Keywords:** tumor growth model, tumor evolution, spatial evolution, ctDNA, tumor DNA shedding

## Abstract

Circulating tumor DNA (ctDNA) monitoring, while sufficiently advanced to reflect tumor evolution in real time and inform on cancer diagnosis, treatment, and prognosis, mainly relies on DNA that originates from cell death via apoptosis or necrosis. In solid tumors, chemotherapy and immune infiltration can induce spatially variable rates of cell death, with the potential to bias and distort the clonal composition of ctDNA. Using a stochastic evolutionary model of boundary-driven growth, we study how elevated cell death on the edge of a tumor can simultaneously impact driver mutation accumulation and the representation of tumor clones and mutation detectability in ctDNA. We describe conditions in which invasive clones end up over-represented in ctDNA, clonal diversity can appear elevated in the blood, and spatial bias in shedding can inflate subclonal variant allele frequencies (VAFs). Additionally, we find that tumors that are mostly quiescent can display similar biases, but are far less detectable, and the extent of perceptible spatial bias strongly depends on sequence detection limits. Overall, we show that spatially structured shedding might cause liquid biopsies to provide highly biased profiles of tumor state. While this may enable more sensitive detection of expanding clones, it could also increase the risk of targeting a subclonal variant for treatment. Our results indicate that the effects and clinical consequences of spatially variable cell death on ctDNA composition present an important area for future work.

## Introduction

A once far-fetched idea that a blood sample can precisely inform on cancer diagnosis, treatment, and prognosis is quickly becoming clinical reality (Wan et al., 2017). This is largely due to advances in the quantification of DNA fragments from cancer cells shed into the bloodstream, known as circulating tumor DNA (ctDNA), which are primarily released from the tumor via apoptosis, necrosis, and active secretion (De Rubis et al., 2019). While tissue biopsies have been a critical component in cancer care, providing a snapshot of the tumor-host microenvironment, they are invasive and repeated biopsies over time to monitor cancer progression and optimize therapies are seldom feasible. Moreover, even when accessible, a single biopsy sample may not represent an entire tumor, which usually displays significant spatial heterogeneity. ctDNA-based “liquid biopsies”, on the other hand, do not have some of these drawbacks and can act as a noninvasive cancer biomarker, allowing analyses of the tumor’s genetic evolution more frequently and comprehensively (Cha et al., 2023; Ulz et al., 2017; Kujala et al., 2022). Two major applications of ctDNA already used in the clinic are for the monitoring of tumor burden before, during, and after treatment, and for the detection of post-treatment relapse (Mattox et al., 2019; Ignatiadis et al., 2021). Liquid biopsies have also shown great promise in predicting relapse, progression free survival, and overall survival across a variety of tumor types and stages (Reinert et al., 2019; Chae et al., 2019; Sanz-Garcia et al., 2022; Cha et al., 2023).

Despite its potential to revolutionize cancer monitoring and treatment, ctDNA can also show poor concordance between blood and tissue, hampering its general clinical utility (Chae et al., 2017; Merker et al., 2018). The main causes for this include access to only minuscule concentrations of ctDNA in a plasma sample, the limits of current sequencing technologies, the confounding effects of non-cancerous mutations and intra-tumor heterogeneity (Jahangiri and Hurst, 2019). While improvements in assay sensitivity and specificity could help to better resolve the ground truth composition of observed ctDNA in a blood sample, we need different methods to better understand and correct possible inaccuracies arising from biased representations of the different tumor clones in ctDNA fragments.

Changes to ctDNA yield and representation of different mutations have been observed before and during chemotherapy, altering the detectability of resistance-causing mutations (Schwaederlé et al., 2017; Ma et al., 2016; Tran et al., 2023). The majority of cfDNA fragments are around 100-160 base pairs long, which is consistent with apoptosis-induced digestion of nuclear DNA into fragments the circumference of a nucleosome (Stroun et al., 2001; Roth et al., 2011; Hu et al., 2021). Elevated apoptosis can increase the amount and clinical detectability of ctDNA in the bloodstream (Marques et al., 2020) and varying apoptosis rates between clones can in theory lead them to become disproportionately represented in the bloodstream (Heitzer et al., 2020). In addition to the intrinsic differences in growth and death rates for different clones, heterogeneity in the tumor microenvironment due to immune infiltration, hypoxia, or treatment onset can also significantly impact rates of apoptosis (Kaufmann and Earnshaw, 2000; Trédan et al., 2007; Murthy et al., 2021; Giordano et al., 2016; Zhou et al., 2006; Marques et al., 2020; Kato et al., 2016; Rostami et al., 2020). These can in turn influence the evolutionary fate of a tumor by altering its local selective pressures and genetic heterogeneity (Meads et al., 2008).

While there are many models studying tumor growth and evolution, the degree to which this underlying genetic distortion between blood and tumor tissue exists, and the evolutionary mechanisms that shape it, are not directly considered, either in models of tumor evolution derived from ctDNA (Abouali et al., 2022), or in clinical studies of ctDNA concordance (Stetson et al., 2019). Recent mathematical models studied how varying the apoptosis rates of tumor cells could influence the time to detection of early-stage tumors (Avanzini et al., 2020) or the effect of differential shedding on the representation of different metastases in ctDNA (Rhrissorrakrai et al., 2023), but ignore the underlying evolutionary process or study neutral, non-spatial evolution. Separately, a model by Fu et al. (2015) showed how reduced chemotherapy exposure in a sanctuary site can promote acquired resistance, but this work did not specifically model the effects on ctDNA genetic distortions.

Here we combine a stochastic model of boundary-driven tumor evolution (Waclaw et al.,2015; Bozic et al., 2019; Chkhaidze et al., 2019; Noble et al., 2022; Lewinsohn et al., 2023) with a model of differential apoptosis and cellular shedding and study the effects of spatially-heterogeneous cellular apoptosis on ctDNA composition and its genetic distortion relative to the tumor tissue. We spatially constrain tumor evolution by assuming that differential drug penetration or immune system infiltration leads to increased cell death and DNA fragment shedding on the edge of the growing tumor. We compare results across a variety of modeling choices, such as differences between quiescent or proliferative tumors, and track the distortion of clones and subclonal mutations in the ctDNA over time.

We find that, as cancers grow and shed DNA into the bloodstream, the clones responsible for expansion into the edge environment are consistently overrepresented in the ctDNA and, in some cases, when progression results in highly heterogeneous tumors, homogeneous regions trapped in the tumor core are underrepresented in the blood. We further show that over-representation of clones from high-shedding tumor regions can lead to differences in the number of detectable subclonal driver mutations, and that the chosen sequencing detection limit can have a complex effect on the extent of the observed genetic differences. We also discuss the potential clinical relevance of distortions in ctDNA genetic variability during clinically significant events, such as the appearance of an expanding subclone or cell turnover-driven increases in clonal diversity.

For liquid biopsy technologies and ctDNA analyses to transform cancer care, from early screening and diagnosis through treatment and long-term follow-up, we need to better understand how to interpret the genetic diversity measured in the blood and how it can be used to inform on the true composition of the tumor tissue. Overall, our results showcase how spatial heterogeneity in apoptosis and cellular shedding across different regions of a tumor can significantly bias the mutational composition of ctDNA and emphasize important directions for further theoretical and clinical investigation into the effect of the microenvironment on ctDNA origin and quantification.

## Methods

### The tumor growth model

While there are many models of tumor growth, to analyze the role of solid tumor spatial structure in shaping the observed variation in ctDNA, we use a model of boundary-driven growth, in which cells on the periphery of a tumor are assumed to experience higher proliferation rates over time, as compared to the tumor core. This type of spatially-restricted growth is usually observed in tissues with weak physical resistance and it can significantly alter tumor evolution by blunting the strength of selection, promoting clonal interference, and increasing mutation burden from the tumor core to its edges (Waclaw et al., 2015; Noble et al ., 2022). Because of its simplicity and well-understood properties, it is an excellent starting point for exploring how spatial variation in apoptosis can impact ctDNA release and can bias the observed genetic differences between blood and main tissue.

In our Eden model, cells grow on a 2D regular lattice and each cell has 8 neighbors (a Moore neighborhood), similar to Waclaw et al. (2015); Chkhaidze et al. (2019); Noble et al. (2022); Lewinsohn et al. (2023). Each simulation begins with a single cell and terminates when the population either goes extinct or reaches a size of 60,000 voxels. In the initial stage of growth, the tumor experiences an environment with death rate *d*_1_. Once the tumor reaches a large enough size (here, a radius of 90 voxels or, equivalently, 3 billion cells) we assume the tumor is detected and treatment can occur that can shrink the initial tumor. After detection, we assume that, due to differential chemotherapy drug penetration or differences in immune infiltration and oxygenation, spatial differences in apoptosis appear between between the tumor core and the edge of the tumor. Specifically, cells in the core, or the sanctuary site (radius *R* ≤ 90), continue to experience death at rate *d*_1_, while on the tumor edge, cells have death rate *d*_2_ ≤ *d*_1_. For the sake of simplicity, we do not model angiogenesis or interactions of cancer cells with other cell types.

This spatial difference in death rates effectively creates a selective barrier for tumor expansion. We consider two parameter regimes: *d*_1_ *< b < d*_2_ and *d*_1_ *< d*_2_ *< b*, which we call “driver-dependent” and “driver-independent” invasion, respectively (**Figure 1**). In the driver-dependent regime, only lineages that have acquired sufficient driver mutations can expand past the core radius *R*, while, with driver-independent invasion, all lineages continue to grow in the presence of the new edge environment. At each time step, a random cell is chosen uniformly from the population, and attempts division with a probability equal to its birth rate *b* * (1 + *s*)^*n*^, where *b* is the baseline birth rate in the population, *s* is the selective advantage of driver mutations, and *n* is the chosen cell’s driver mutation count. If the cell attempts division, it places a daughter cell in a randomly-chosen empty site in its Moore neighborhood. If the cell is completely surrounded, it cannot divide. Upon division, we assume that the daughter cell acquires a Poisson-distributed number of additional driver mutations, with rate *μ*. We assume each mutation appears only once (infinite site assumption). After attempting division, the chosen cell is removed from the population with probability equal to its death rate *d*_*i*_, where *i* ∈ 1, 2 indicates which region of the tumor the cell inhabits.

**Figure 1:**
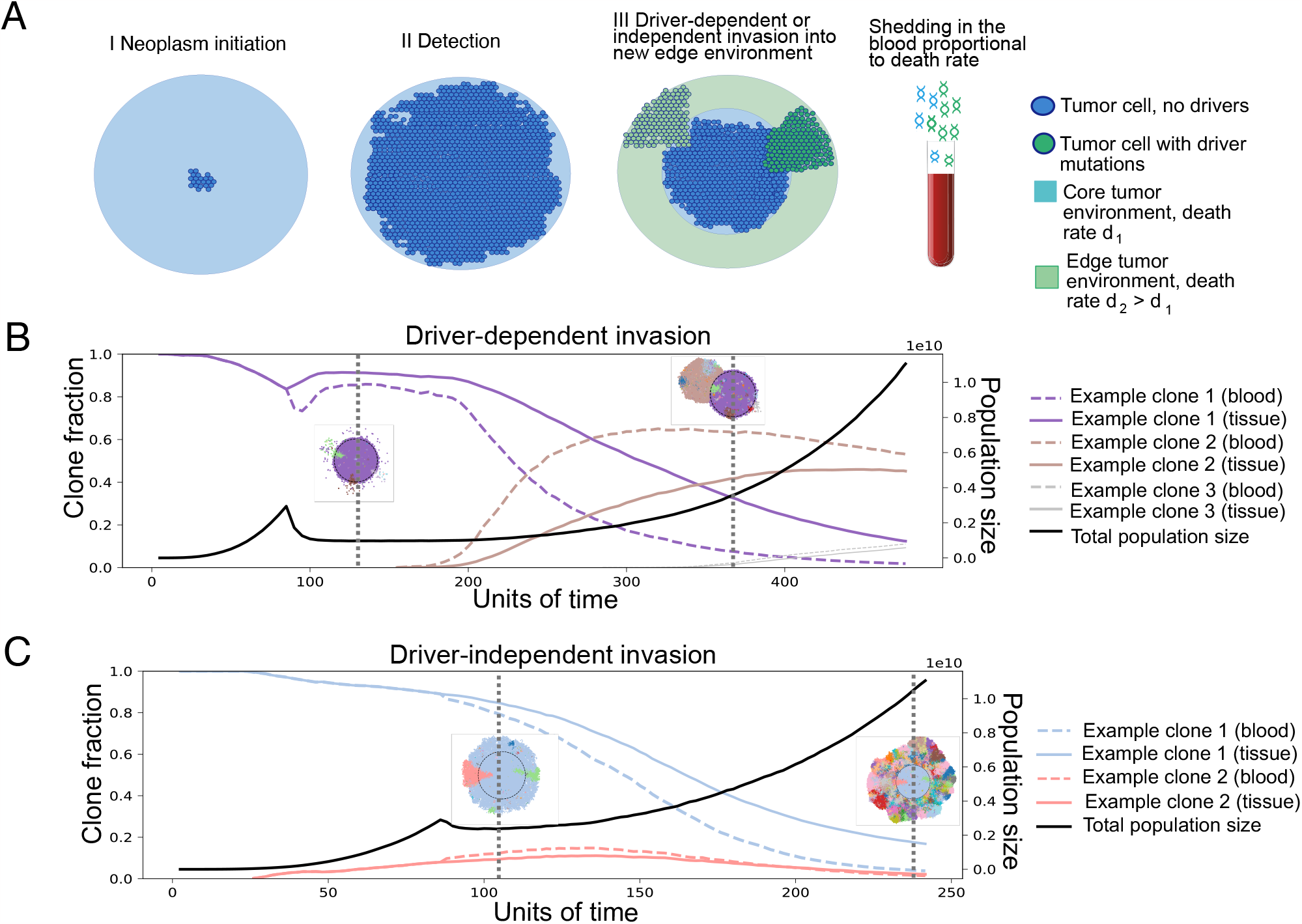
**A**. Illustration of the model. Tumors grow to a clinically detectable size (a 2D cross-section of a 3 billion cell tumor), and are then partially exposed to a new environment, where the cells die with rate *d*_2_. The growth rate in the new environment determines the invasive potential of a clone. If the death rate *d*_2_ is higher than the initial birth rate, only clones with mutations increasing the growth rate to a positive number can grow in the new environment, so invasion is driver-dependent. Otherwise, it is driver-independent. Tumor growth can be proliferative or quiescent. In the former, cells divide when they have an empty neighbor on the lattice and die at a rate independent of their neighbors. In the latter, cells also divide when they have an empty neighbor on the lattice, however cell death also requires empty neighbors. The shedding rate of DNA into the blood is assumed to be proportionate to the death rate. **B**. Example trajectories, driverdependent invasion. Trajectories of clone fractions and total population size for driver dependent invasion, with visualizations of the 2D tumor at selected timepoints. Each color corresponds to a unique clone, also shown in the trajectory plot. **C**. Example trajectories, driver-independent invasion. Trajectories of clone fractions and total population size for driver independent invasion, with visualizations of the 2D tumor at selected timepoints. For both cases, *μ* = 0.001, *s* = 0.1, *d*_1_ = 0.1, *b* = 0.7. For driver-dependent invasion, *d*_2_ = 0.9. For driver independent invasion, *d*_2_ = 0.69.

We also analyze a version of the main model where cells do not die if they are fully surrounded, so that the tumor core remains in a quiescent state and where selection acts by reducing the apoptosis rate rather than increasing birth rate, so that *d* ← *d* * (1 − *s*).

#### Parameter Choices

To significantly save on simulation time and memory, we assume a Poisson distributed driver mutation rate of *μ* = 0.001, roughly 100 times the estimated empirical rate, which we denote by *μ*_*real*_ = 1*e* − 5, as in Bozic et al. (2019). We also simulate the tumors in 2D, so that the spatial heterogeneity reflects that of a cross section of a much larger 3D tumor, a rationale used in Noble et al. (2022) for similar 2D spatial models. Each 2D voxel then represents 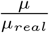 identical cells. For a simulation with *m* voxels, we roughly approximate the 3D tumor size, *N*, to be that of a sphere, with a cross section equal in area to the number of 2D cells, such that 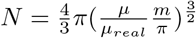. We further choose a sanctuary site radius, *R*, ranging from 20 to 60 voxels. Assuming 20*μm* diameter tumor cells, and 100 cells per 2D voxel, this *R* would correspond to an equivalent tumor with a radius of 0.4 to 1.2cm and approximately 1000 to 20,000 cells, representing a cross section of a 3D tumor of roughly 30 million 1 billion cells (Del Monte, 2009). We simulate tumors until they expand well beyond the core sanctuary site and stop the simulations when tumors reach a size of 60,000 voxels, corresponding to a tumor size of approximately 10 billion cells or a radius of 2.5cm. Without loss of generality, throughout what follows, we also assume a constant selective benefit for driver mutations, *s* = 0.1.

**Table 1:** Main parameters used in the model.

|  |  |
| --- | --- |
| $N$ | Final tumor size |
| $R$ | Core / sanctuary site radius |
| $b$ | Initial cell birth rate |
| $d_1$ | Cell death rate in the tumor core |
| $d_2$ | Cell death rate in the tumor edge |
| $s$ | Driver mutation fitness advantage |
| $\mu$ | Poisson-distributed driver mutation rate |

### Modeling clone fractions and variant allele frequencies (VAFs) in ctDNA

To compute clone frequencies in the ctDNA, let *N*_*ij*_ be the number of cells of clone *i* from region *j*, with corresponding death rate *d*_*j*_. We assume that shedding into the blood is proportional to the death rate of a tumor region (Avanzini et al., 2020), i.e. the fraction of a tumor clone in the ctDNA population at time *t* can be computed as a weighted average over the frequency of the clone in each region,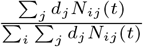.

While this represents the clone’s fraction of the tumor population, to test the effect of clone fraction distortion on mutation detection, we also estimate clinically realistic VAFs in the blood, which also contains DNA fragments from healthy tissue. To do this, we compute the frequencies of each driver mutation belonging to each clone and then estimate the fraction of the total number of fragments that originate from the tumor (the tumor fraction). At the point of diagnosis, Phallen et al. found that the mean tumor fraction in the bloodstream for stage I and II breast, lung, ovarian, and colorectal tumors was 1% (Phallen et al., 2017). We calibrate the simulated tumor fraction by assuming this is the fraction for proliferative tumors at the point of detection, assumed to occur at 3 billion cells, with initial death rate of *d*_1_ = 0.1.

To estimate a shedding probability, we adapt a formula from Avanzini et al. (2020). Assuming an exponentially growing tumor with a constant growth rate, the formula computes the number of fragments shed into the bloodstream as a Poisson-distributed random variable, with mean 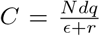, where *N, d, q,ϵ* and *r* are the number of cells, death rate, shedding rate, decay rate, and growth rate respectively. We estimate C using the Phallen data set, which found the median DNA concentration in plasma to be 29 ng/ml. Repeating a calculation from their paper, a haploid genome weighs roughly 0.0033ng, suggesting that there are 8788 haploid genome equivalents (HGEs) in 1ml of plasma. With 5L total blood volume in the human body and 55% plasma, we can therefore estimate *C* to be 5000 × 0.55 × 8788 × 0.01 = 241, 670. While the formula depends on *r* (the tumor birth rate can in fact slightly alter the total ctDNA molecules present in a blood draw), the tumor population changes on the order of days, while DNA decays in the blood with a half life of about 30 minutes (Sanz-Garcia et al., 2022). This implies *ϵ* = 48 ln 2 ≈ 33.3, while *r <* 1. In a spatial setting, the effective growth rate is even lower, because cells do not divide when surrounded, so we assume *r* ≈ 0. Setting 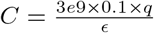, we estimate *q* ≈ 0.026.

The mean number of tumor fragments at other time points is then computed as 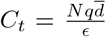,where 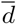the mean death rate of the whole tumor. For a mutation *m* with tissue frequency *f*_*m*_ and overall death rate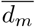, we write the total number of fragments with that mutation as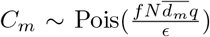. For a 15ml blood draw (0.3% of the total supply), we scale the mean number of fragments by 0.003. Let 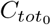fragments in a 15ml blood draw, at the point of detection. Then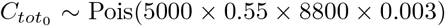.We assume the mean number of non-tumor fragments remains constant at 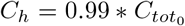.If we assumeall cells are diploid, each mutation appears on a single chromosome copy and we ignore the possibility of recurrent mutation or subsequent allelic gain or loss, we can write the expression for the spatially biased VAF of a specific mutation in the blood as 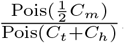. To analyze the effect of spatially correlated death rates on the detection of tumor mutations, we compute both spatially biased and unbiased VAFs by using the mean death rate of the specific mutation 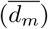 for the former, and the mean death rate of the entire tumor (replace 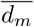with 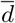in the expression for *C*_*m*_) for the latter.

#### Inverse Simpson diversity as a measure of intratumor heterogeneity, ITH

Since an important goal of this work is understanding how ctDNA data collected from the blood may distort estimates of clonal heterogeneity present in the main solid tumor, we use the inverse Simpson diversity index to quantify and compare heterogeneity estimates from blood and tissue sequences. The inverse Simpson diversity index is a classic diversity measure employed in many previous studies of population diversity which takes into account the number of lineages present, as well as the relative abundance of each (Buckland et al., 2005; Noble et al., 2022). For a set of clone fractions *f*_1_, …, *f*_*N*_, with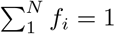, it is defined as 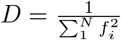.

## Results

### Spatial differences in apoptosis and shedding can bias clone fractions in ctDNA

To study how the spatial structure of a solid tumor, through spatial heterogeneity in apoptosis, can bias the observed ctDNA in blood draws, we first analyze the difference in the clonal fractions between blood and tumor tissue. In **Figure 2**, we compare results for proliferative versus quiescent cell models, small versus large sanctuary sites and driver-dependent versus driver-independent invasion. Across all modeling scenarios, **Figure 2** shows that new clones on the expanding front tend to be over-emphasized in the ctDNA, while older clones, trapped in the tumor sanctuary, tend to be under-represented. The magnitude of the differences in clonal fraction and their likelihood to impact clinical detectability depend on the accumulated clonal diversity on the edge of the tumor, mediated by the edge environmental effects.

**Figure 2:**
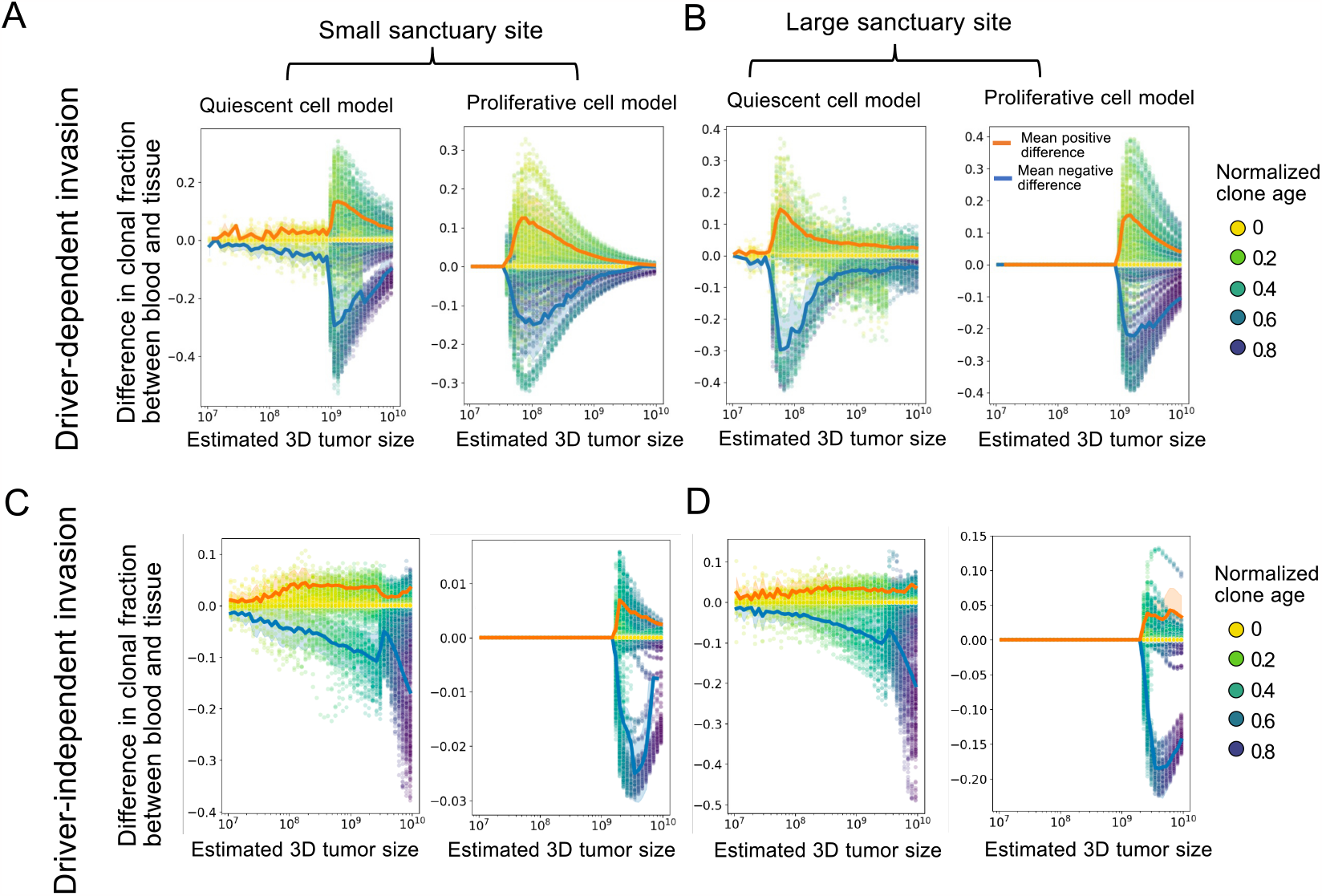
Clone fraction differences between blood and tissue: **(A-D)** Each plot shows the results of 50 simulation runs, where each point represents the difference between clonal frequencies estimated from the blood versus those present in the tumor, for a single clone, with the color showing the age of the clone relative to the total simulation time. Tumors were grown from a single cell until reaching a 2D cross-section of a 10 billion cell tumor. For all simulations, *μ* = 0.001, *s* = 0.1, *d*_1_ = 0.1, *b* = 0.7. For driver-dependent invasion, *d*_2_ = 0.9. For driver independent invasion, *d*_2_ = 0.69. The orange and blue lines show the average positive and negative clone fraction difference, respectively. Only clones comprising at least 10% of the tumor were included in the average. Shading is ±1 s.d. We show the same plots over normalized time in **Supplementary Figure S2**.

In the driver-dependent case (**Figures 2A** and **B**), the few driver clones able to invade the new environment experience a higher death rate during expansion on the edge and end up over-represented in the blood, making the absolute difference between the blood and tissue clone fractions substantial. The maximum difference between the two occurs in the limiting case of a single clone, originating on the expanding front and growing without competition in the new edge environment. For proliferative tumors, we can write an upper bound for this clone fraction difference. If we assume the tumor initiates with death rate *d*_1_ and grows to a constant size *S*, after which a single invasive subclone grows to size *x*, experiencing death rate *d*_2_, the difference in the expected clone fraction can be written as

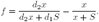

It is easy to show that the maximum value of *f* is 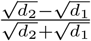, which occurs when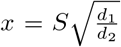. We plot the maximum possible clone fraction difference for all *d*_1_ and *d*_2_ in **Supplementary Figure S1** and show that despite the apparently high choice of *d*_2_ in some of our simulations, large differences in estimated clonal frequencies can occur with very small absolute death rates. In line with the prediction that the peak clone fraction difference does not depend on region size, simulations also show that, for driver-dependent invasion, the size of the tumor sanctuary does not greatly impact the distribution of clonal fraction differences (**Figures 2A** and **B**).

The sanctuary size does affect the results for proliferative driver-independent tumors, which show very little difference between the ctDNA and main tissue, when the sanctuary site is small (**Figure 2C**). This is because early clones from the small sanctuary region can invade the edge environment before the appearance and spread of later clones, and are therefore represented throughout all tumor regions that differentially shed into the blood. This effect is still present with a larger sanctuary site, since the observed minimum clonal fraction difference is still much smaller than the corresponding one in the driver-dependent case (compare **Figures 2B** and **D**).

For quiescent tumors, ctDNA can only come from the shedding of cells on the expanding front, which is determined by the total size of the tumor prior to detection, and the sanctuary size again has little effect on the observed differences (**Figures 2B** and **D**). Despite this, the magnitude of the differences in death rates are comparable to proliferative tumors. However, we notice that quiescent tumors distort clone fractions across all population sizes and time points, due to the additional spatial bias in death rate. One thing to note is that, while here we assume that differences in shedding are caused by spatial heterogeneity in death rates, we expect results to be similar in any extension of the model in which clones are weighted differently in the ctDNA than the tissue, for example, with differential access to the bloodstream based on proximity to blood vessels or via a model of active secretion. Additionally, we find that the version of the model where driver mutations reduce death rate, akin to apoptosis resistance, results in similar clone fraction distortions (**Supplementary Figure S3**).

### Differential shedding can make us overestimate the true intra-tumor heterogeneity

In **Figure 3** we use the inverse Simpson diversity index across normalized time points as a proxy for ITH in the ctDNA and in the tissue, over the course of tumor progression. We find that driver-independent tumors with a large sanctuary site consistently show a large difference between blood and tissue ITH (**Figure 3D**), while tumors with a small sanctuary site do not show any difference. This is a consequence of the clone fraction differences observed in **Figure 2**, which, for proliferative tumors, vanish once the sanctuary site is too small. Also consistent with **Figure 2**, quiescent driver-independent tumors show elevated ITH for both sanctuary sizes (**Supplementary Figure S4**). As expected, driver-dependent tumor growth is driven by very few clones following detection, which results in much lower overall clonal diversity (**Figures 3A, B** and **Supplementary Figure S4**).

**Figure 3:**
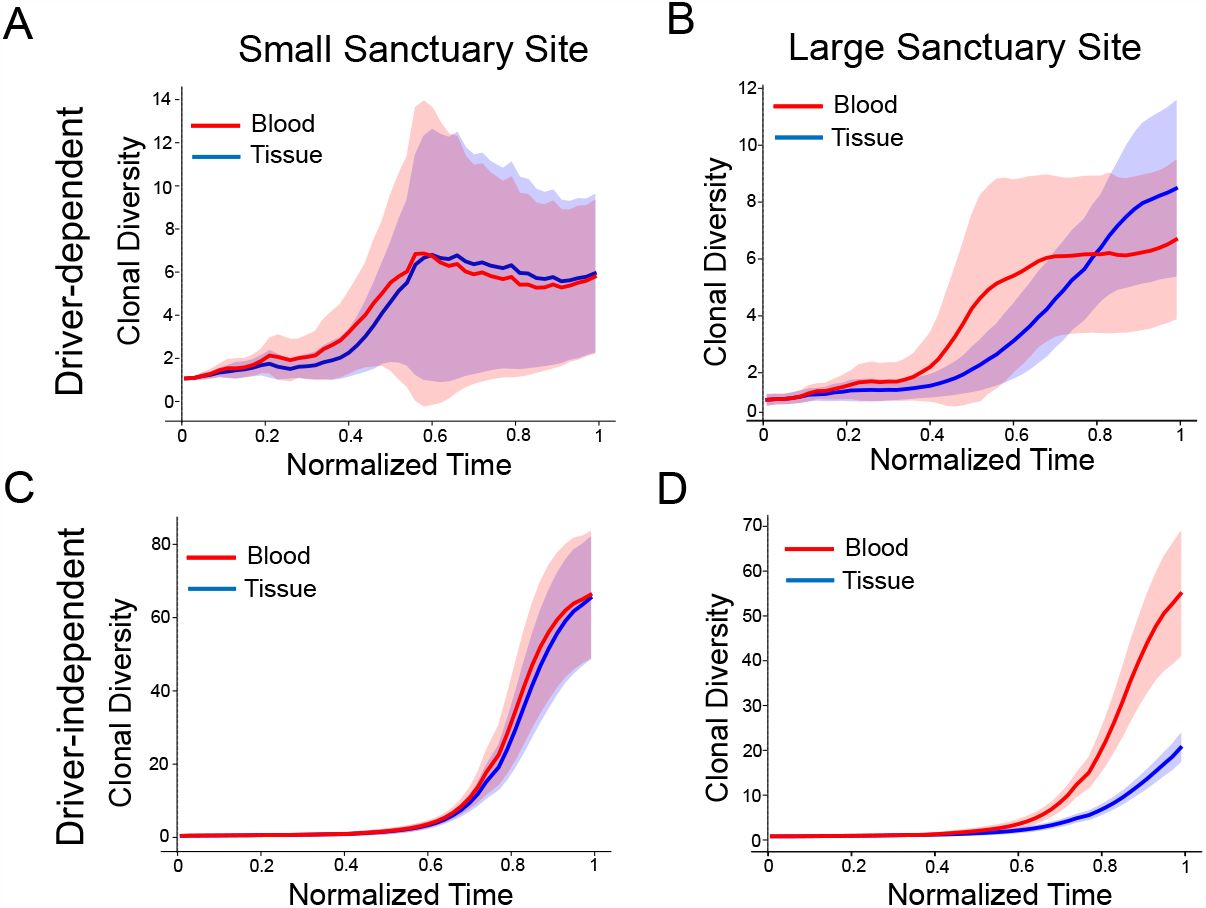
Discrepancies between blood and tissue clonal diversity. The subplots show the inverse Simpson diversity index of the clonal frequencies in the blood and tissue for each clone in 50 simulated tumors. Timepoints are normalized by run and then binned and down-sampled. Tumors were grown from a single cell until reaching a 2D cross-section of a 10 billion cell tumor. For all simulations, *μ* = 0.001, *s* = 0.1, *d*_1_ = 0.1, *b* = 0.7. For driver-dependent regrowth, *d*_2_ = 0.9. For driver independent regrowth, *d*_2_ = 0.69. Shading represents ±1 s.d. The figure shows results for proliferative tumors only. For all scenarios, see **Supplementary Figure S4**.

### The effect of sequencing detection limits and sanctuary site size on observed VAFs in the blood

We next analyze how biased clonal fractions in the blood translate into biased observed VAFs, under various sequencing detection limits. In **Figure 4**, we consider sequence detection limits of 10^*−*3^ and 10^*−*2^, which are often utilized for panel-based assays optimized for MRD detection (Chin et al., 2019). As expected, a higher sequence detection limit of 10^*−*2^ diminishes the number of detected drivers (VAF exceeds the detection limit) and increases the tumor size at which the first mutations are detected, compared to a limit of 10^*−*3^ (**Figure 4A**). This effect is more pronounced in quiescent tumors than proliferative ones. While driver-independent tumors produce many more mutations, responsible for the higher ITH shown in (**Figure 3**), they are nonetheless very low-frequency and so the number of mutations above a 10^*−*2^ threshold is comparable to that of driver-independent tumors. Most mutations evade detection entirely, as the overall percentage of driver mutations detected at any point is below 10% for all scenarios (**Supplementary Figure S5 C-D**).

**Figure 4:**
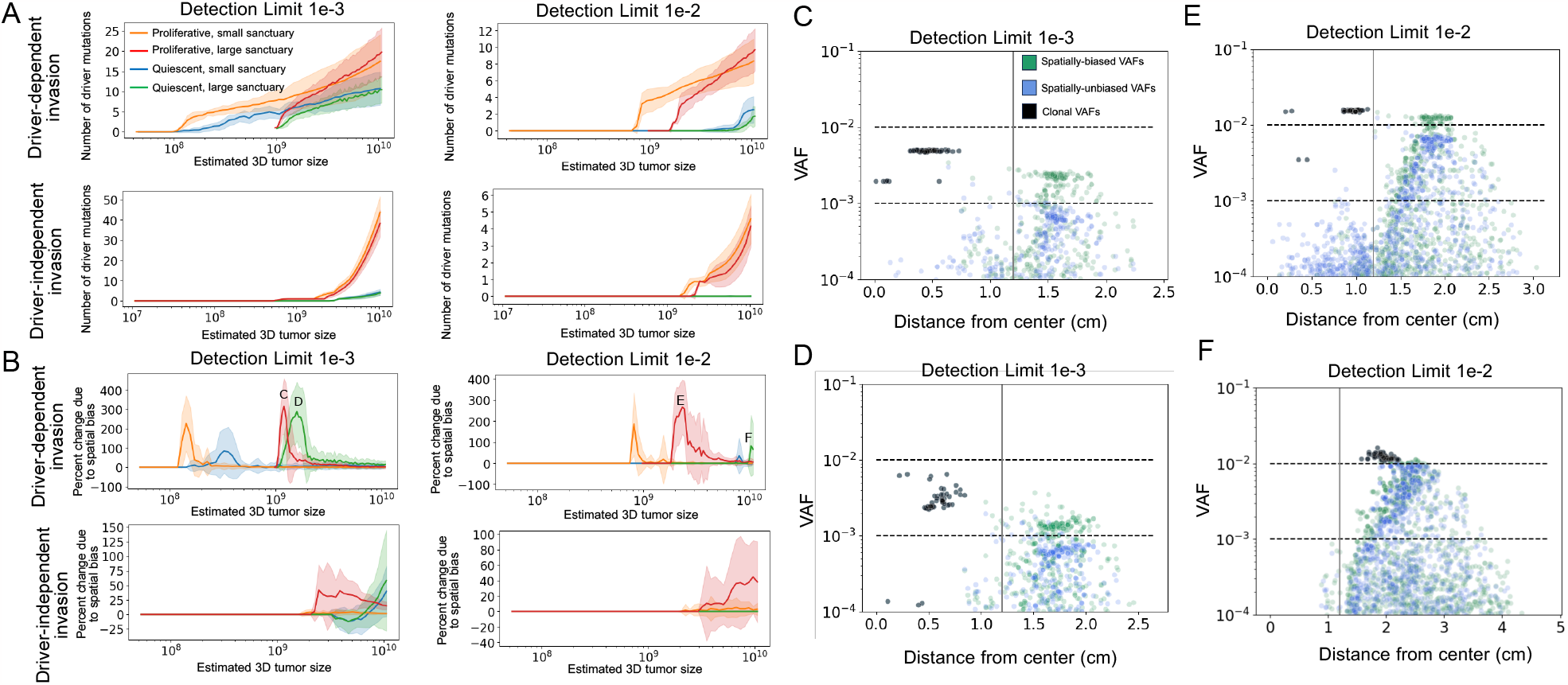
Influence of spatial bias on limits of detection. **A**. Plots of the number of detectable driver mutations starting from the point of relapse for minimum detection frequencies of 1e-3 and 1e-2 for proliferative and quiescent tumors relapsing at ∼10^8^ and ∼10^9^ cells. Mutations were detectable if the estimated VAF exceeded the detection limit. VAFs were estimated based on a tumor fraction of 1% for a 3 billion-cell tumor with death rate of 0.1 (see Methods). **B**. Percent change in number of detectable drivers when the VAFs in **A** are compared to VAFs computed assuming the tumor sheds all clones at the same rate for the same detection limits (see Methods). **C-F**. Scatter plots of mean spatially biased VAFs (green) and unbiased VAFs (blue) at the size where the average spatial bias over all replicates is maximal (marked with the corresponding letter in **B**). Each plot shows all mutations for 50 replicates of the corresponding simulation scenario. The x-axis is the mean distance of the mutation from the tumor’s center. Black points are clonal mutations, which show perfect overlap between the blood and tissue. The vertical line marks the end of the sanctuary region.

In **Figure 4B**, we compare the percent change in number of detectable drivers when the simulated VAFs are compared to VAFs from a spatially uniform null model, computed assuming the tumor sheds all clones at the same rate. We show that spatial tumor heterogeneity can greatly affect the number of detectable driver mutations in the blood, and sequencing detection limits can further alter the extent of this bias, with the timing and magnitude of difference spikes further dependent on the detection limit of the sequencing technology.

Because clonal VAFs cannot change due to shedding differences, the effect depends entirely on the detection limit relative to subclonal VAFs. We see that spatial bias in proliferative driver-dependent tumors increases when the detection limit is raised, but quiescent spatial bias either decreases in magnitude and appears at a larger tumor size, or disappears all together. We show the percent spatial bias over normalized time in **Supplementary Figure S5 B**.

In **Figures 4C-F**, we examine the dependency of spatial bias on detection limit by plotting the frequency versus the mean tumor radius of every mutation present in 50 simulation runs at the point of maximal spatial bias (the labeled peaks in **Figure 4B**). Plots corresponding to the peaks of the other scenarios are shown in **Supplementary Figure S6**. We observe a cluster of clonal mutations in the core of the tumor (colored black), which are equally represented in the blood and tissue. Due to boundary-driven growth, subclonal mutations accumulate more on the edge of the tumor and tend to remain there across generations, increasing the frequency of mutations further from the core. Because the mutations also shed at higher rates, filtering for larger mutations can increase bias, but will decrease it once the majority of detectable VAFs are clonal (**Figure 4F**). Of clinical relevance is the case where subclonal variants are exaggerated to near-clonal frequencies, which occurs in the driver-dependent case (**Figure 4C-F**). This showcases the benefits and risks of distorted ctDNA: while exaggerated subclonal mutations would provide more biomarkers to aid in detecting recurrence, they would make poor targets for treatment.

## Discussion

As cancers grow, they slough off cells and DNA from apoptotic or necrotic cancer cells, which enter the bloodstream. Through the use of technologies such as next-generation sequencing, these fragments of DNA can reveal a wealth of information about cancer, without the need for invasive surgical biopsies. Here we explore how boundary-driven tumor growth and spatial heterogeneity in cellular death rates impact both the clonal evolution of the tumor, and its representation in ctDNA. We find that the appearance of genetic distortions between blood and tissue ultimately depends on whether the tumor’s genetic heterogeneity varies with respect to rates of apoptosis and ctDNA shedding, which themselves can vary between tumors or over time, for a single tumor. When there is a strong correlation, such as when a change in cellular death rate occurs in direction of tumor growth, ctDNA can drastically bias which clones are observed and can lead to biased estimates of intratumor heterogeneity (ITH).

In the driver-dependent case and, to a lesser extent, the driver-independent case explored here, this bias can be beneficial, by increasing the visibility of and sensitivity for the particular mutations responsible for tumor progression. Spatial differences in cell death rates could also lead to subclonal mutations appearing at clonal frequencies in ctDNA, thus increasing the likelihood that they are mistaken for clonal mutations and chosen as therapeutic targets (**Figure 4**). Our results agree with findings that quiescent tumors may be difficult to detect in the bloodstream (**Figure 4A**), and further suggest that any detectable ctDNA is likely to dramatically under-represent some tumor regions with reduced shedding (**Figure 2**). One possibility is that a lesion with a quiescent interior could be nearly undetectable and suddenly begin to shed appreciably due to a clonal expansion. Because of the extremely biased location of shedding in quiescent tumors, the overall size should not be assumed to correlate well with ctDNA yield. The potential for exaggerated observed heterogeneity in the blood relative to the tissue for tumors experiencing high apoptosis on the expanding front suggests that low-frequency clones, with a high probability to go undetected in a tissue sample, could be better captured in the blood and provide an early indicator of heterogeneous growth. At the same time, when clinical studies find greater heterogeneity in blood than in tissue samples, this is usually mainly attributed to missed heterogeneity in the tissue sample. However, localized high death rates could generate more mutations and at the same time enrich these in ctDNA, through increased shedding. This both poses a potential confounding factor for assessing tumor mutational burden from ctDNA, while simultaneously supporting the potential of blood-based diagnostics to be a more sensitive indicator of changing levels of heterogeneity than tissue biopsies. Recent work has found that in contrast to high tissue mutational burden, which may indicate high neoantigen load and better overall survival, high blood mutational burden may better reflect overall ITH and therefore indicate poor overall survival (Fridland et al., 2021). High heterogeneity correlated to high-shedding regions could contribute to this discordance.

This general principle that genetic distortion between blood and tissue is a function of clonal dynamics is not limited to spatial heterogeneity in intrinsic death rates, and could also arise as the result of differential access to blood vessels or nutrients. Further specific scenarios can be theoretically and clinically explored, such as local metastasis of a primary breast tumor to the lymph nodes, or the microinvasion of a colorectal tumor into the subserosal tissue, particularly during neoadjuvant treatment when the tumor faces novel selective pressure. In both of these cases, there is recent evidence that ctDNA shedding can vary as a function of spatial location. Clonal replacement during treatment for early stage breast tumors is also well documented, and a small study of early stage breast cancer patients discovered mutations private to clones that invaded the lymph nodes. In one patient, as an example of subclone over-representation, these mutations comprised the majority of detected ctDNA (Bredno et al., 2020; Caswell-Jin et al., 2019; Barry et al., 2018).

While our simulations consider only a single form of spatial growth and do not incorporate a fully realistic downstream analysis of ctDNA, here we nonetheless show that even a simple model of spatially heterogeneous tumor growth and shedding can showcase how blood sample data may not represent the tissue accurately, depending on the evolutionary processes shaping the tumor around the time of a blood draw. Further biases as a result of low tumor fraction in cfDNA, copy number variation, germline mutations, hematopoetic mutations, and heterogeneity absent from small tissue samples introduce significant additional complexity that we ignore here (Kammesheidt et al., 2018; Chan et al., 2020). Future directions include incorporating a spatial model of blood vessel distribution that impacts drug delivery, oxygenation, and the resulting apoptosis and shedding rates. Rather than modeling changes to overall clone frequencies under an infinite sites assumption, incorporating a specific resistance model would further allow predictions of the detectability of specific drivers. Here we assume that changes to birth and death rates happen incrementally through a series of point mutations, while specific models of chemotherapy resistance or immune escape may have a different effect on growth rates and the resulting shedding. Because the expanding clones in our model continue to experience high apoptosis, our results would best apply when apoptosis reduction is absent or only partial in the resistant population, such as in apoptosis-induced compensatory proliferation (AICP) (Friedman, 2016).

A further area of study is using model insights to correct for the observed bias between ctDNA and tissue genetics. The work here reveals some of the circumstances in which we would expect such a bias to manifest and the mechanisms by which it would occur, but systematically inverting that bias to reconstruct with maximum fidelity the clonal composition of the tumor from the blood data will require further work. For example, some important applications of tumor genome samples to clonal lineage tracing (“tumor phylogenetics”) depend on accurate quantification of allele frequencies, and extending such methods to use blood data productively will require ways to not only identify, but also correct for these biases. It will be important to characterize the circumstances under which this problem is invertible and what additional data might be needed.

At a basic level, ctDNA can reveal information about the likely presence and burden of cancer within the body. To make full use of this new technology, further work is needed to understand all of the ways that ctDNA can provide a distorted mirror of the main tissue, how tumor evolution shapes these biases and how to correct for them.

## Code and data availability

Code and raw data used to generate all results and figures for this paper can be found at https://github.com/trachman1/lattice-tumor-ctdna.

## Acknowledgments

We gratefully acknowledge support from the NIH National Institute of General Medical Sciences (award no. R35GM147445) and from the NIH T32 training grant (no. T32 EB009403). Research reported in this publication was also supported by the National Human Genome Research Institute of the National Institutes of Health under award number R01HG010589. This research was done using resources provided by the Open Science Grid, which is supported by the National Science Foundation award 1148698, and the U.S. Department of Energy’s Office of Science. The content is solely the responsibility of the authors and does not necessarily represent the official views of the National Institutes of Health.

## Supplementary Material: Supplementary Figures

**Figure S1.**
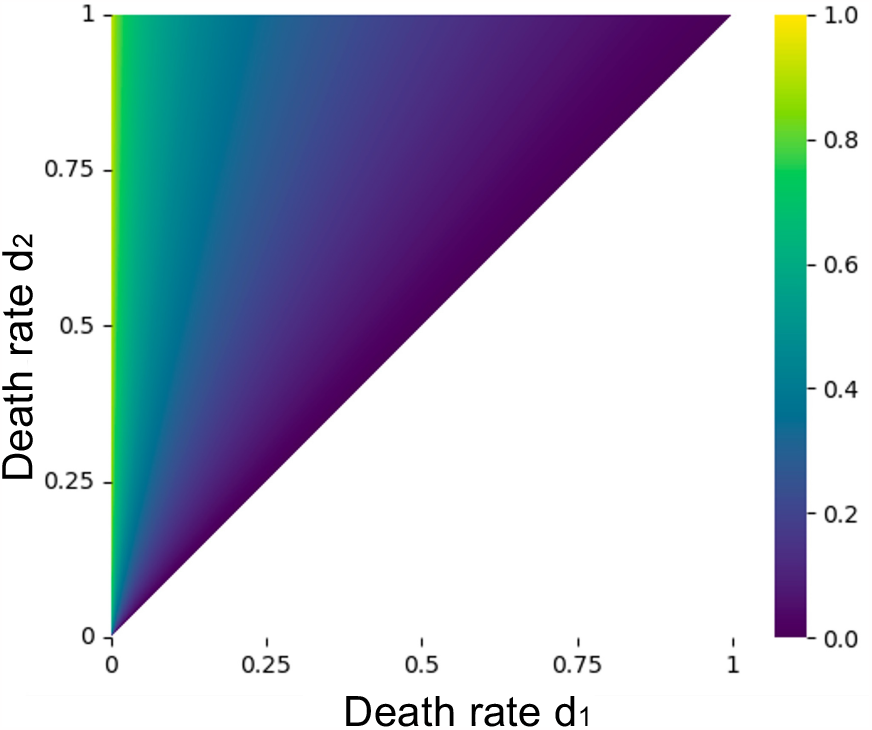
A heatmap showing the maximum clone fraction difference possible for proliferative tumors with respect to all values of *d*_1_ and *d*_2_.

**Figure S2.**
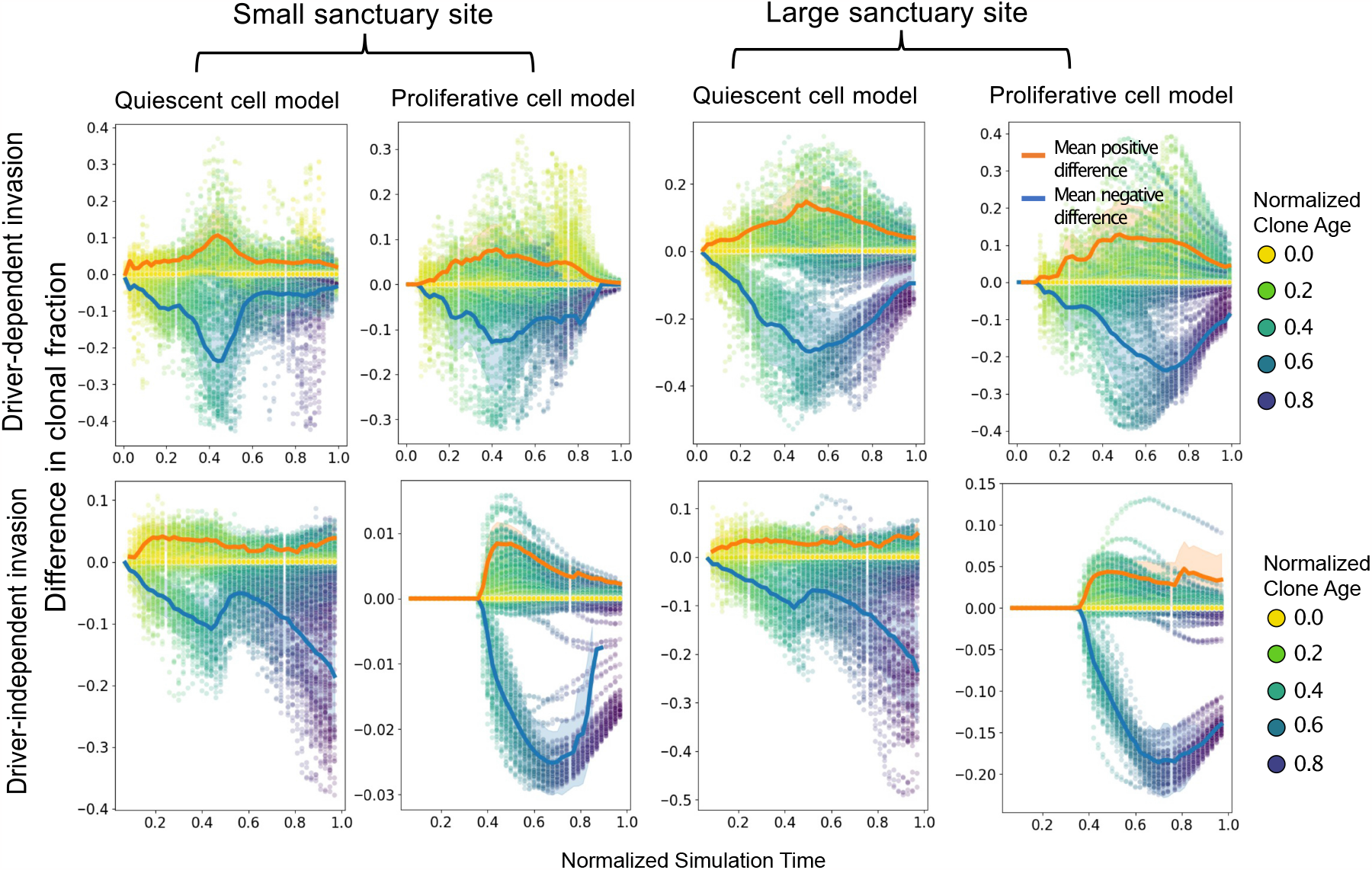
Clone fraction differences between blood and tissue over normalized time: **(A-D)** Each plot shows the results of 50 simulation runs, where each point represents the difference between clonal frequencies estimated from the blood versus those present in the tumor, for a single clone, with the color showing the age of the clone relative to the total simulation time. Tumors were grown from a single cell until reaching a 2D cross-section of a 10 billion cell tumor. Because mutation accumulation is random, we used down-sampled, normalized time points to plot each simulation run within a similar time frame. For all simulations, *μ* = 0.001, *s* = 0.1, *d*_1_ = 0.1, *b* = 0.7. For driver-dependent relapse, *d*_2_ = 0.9. For driver independent invasion, *d*_2_ = 0.69. The orange and blue lines show the average positive and negative clone fraction difference, respectively. Only clones comprising at least 10% of the tumor were included in the average. Shading is ±1 s.d.

**Figure S3.**
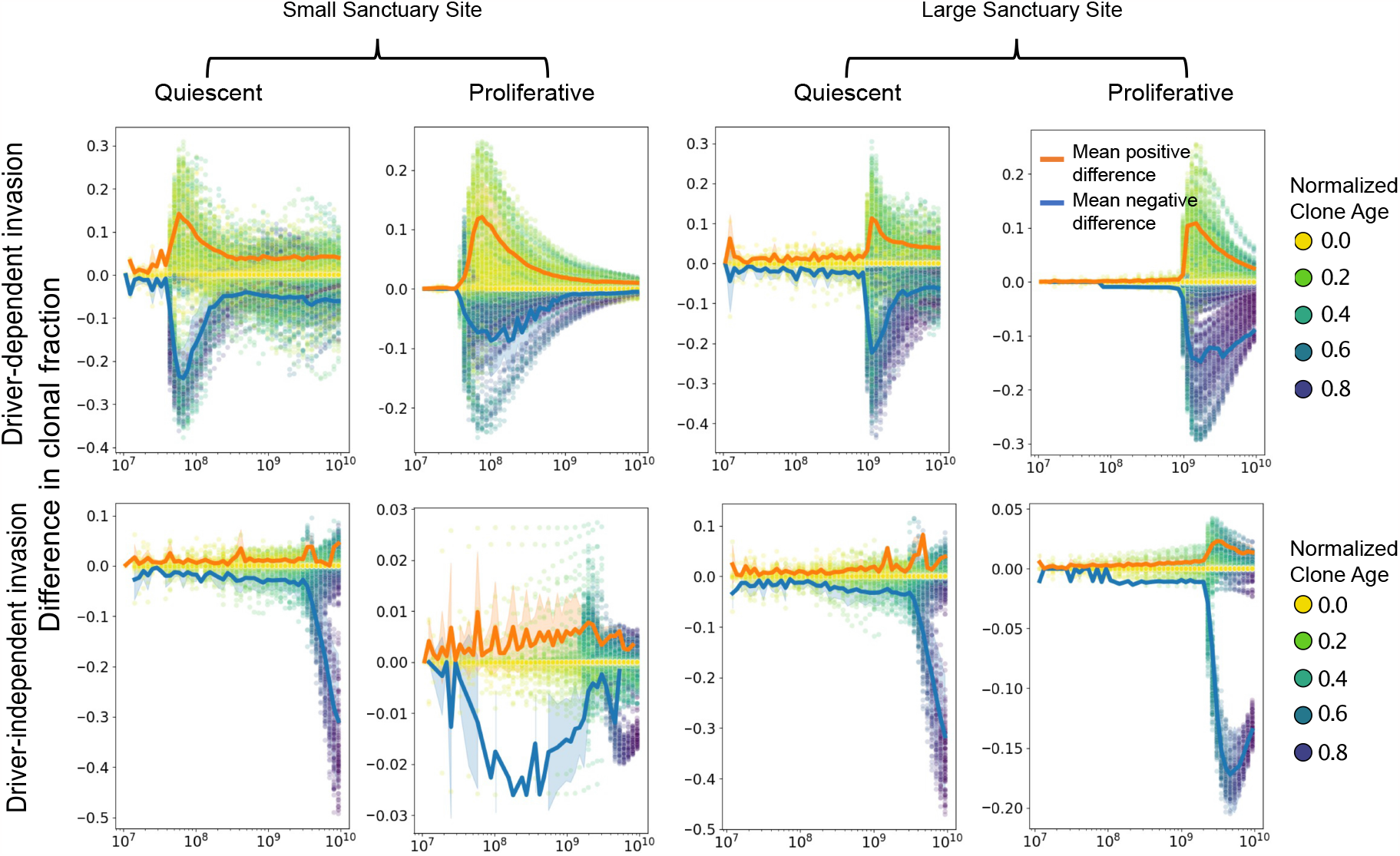
Clone fraction differences between blood and tissue with selection acting on death: **(A-D)** Each plot shows the results of 50 simulation runs, where each point represents the difference between clonal frequencies estimated from the blood versus those present in the tumor for a single clone, with the color showing the age of the clone relative to the total simulation time. Tumors were grown from a single cell until reaching a 2D cross-section of a 10 billion cell tumor. For all simulations, *μ* = 0.001, *s* = 0.1, *d*_1_ = 0.1, *b* = 0.7. For driver-dependent relapse, *d*_2_ = 0.9. For driver independent invasion, *d*_2_ = 0.69. The orange and blue lines show the average positive and negative clone fraction difference, respectively. Only clones comprising at least 10% of the tumor were included in the average. Shading is ±1 s.d.

**Figure S4.**
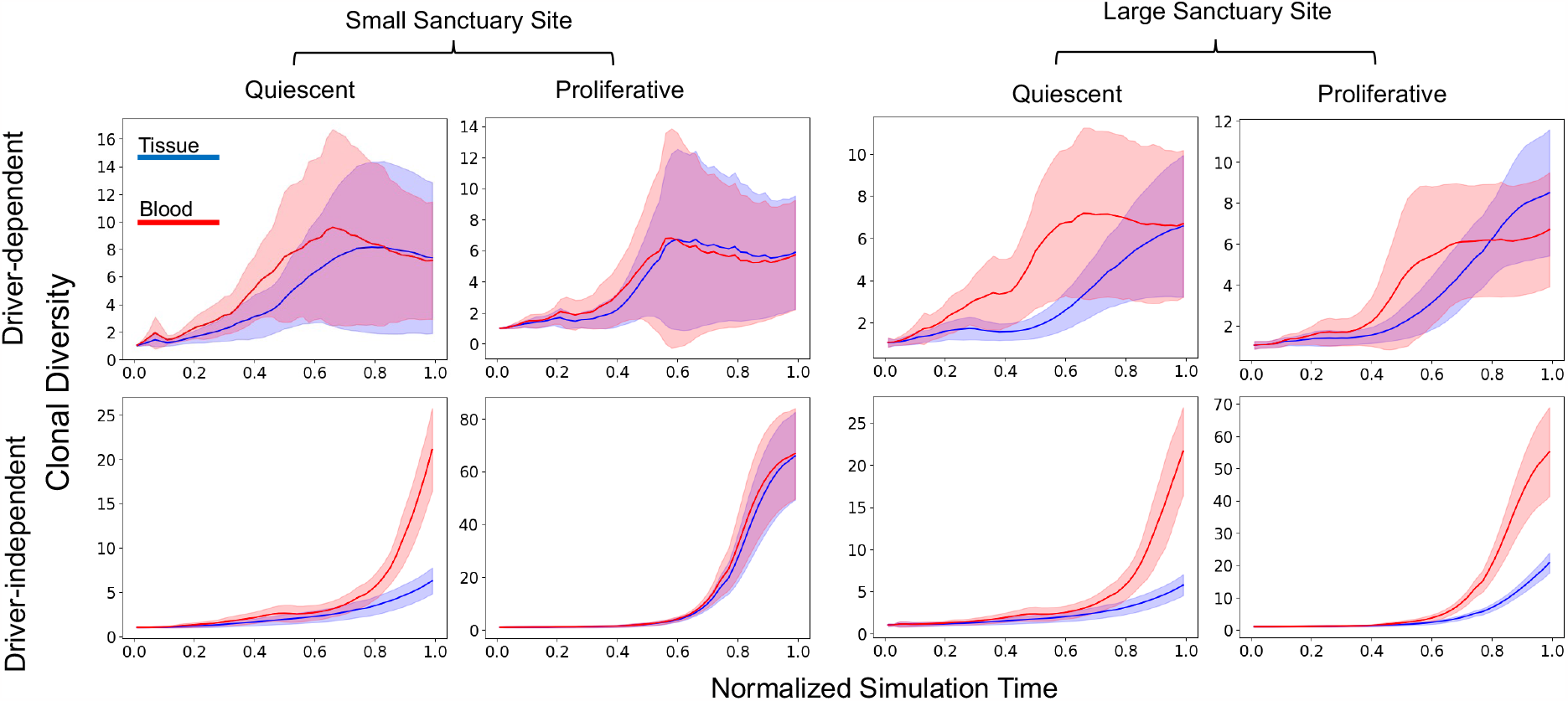
Discrepancies between blood and tissue clonal diversity: Inverse Simpson index of clone frequencies in blood and tissue for each clone in 50 simulated tumors at simulation timepoints normalized by run and then binned and down-sampled. Tumors were grown from a single cell until reaching a 2D cross-section of a 10 billion cell tumor. For all simulations, *μ* = 0.001, *s* = 0.1, *d*_1_ = 0.1, *b* = 0.7. For driver-dependent regrowth, *d*_2_ = 0.9. For driver independent regrowth, *d*_2_ = 0.69. Shading represents ±1 s.d.

**Figure S5.**
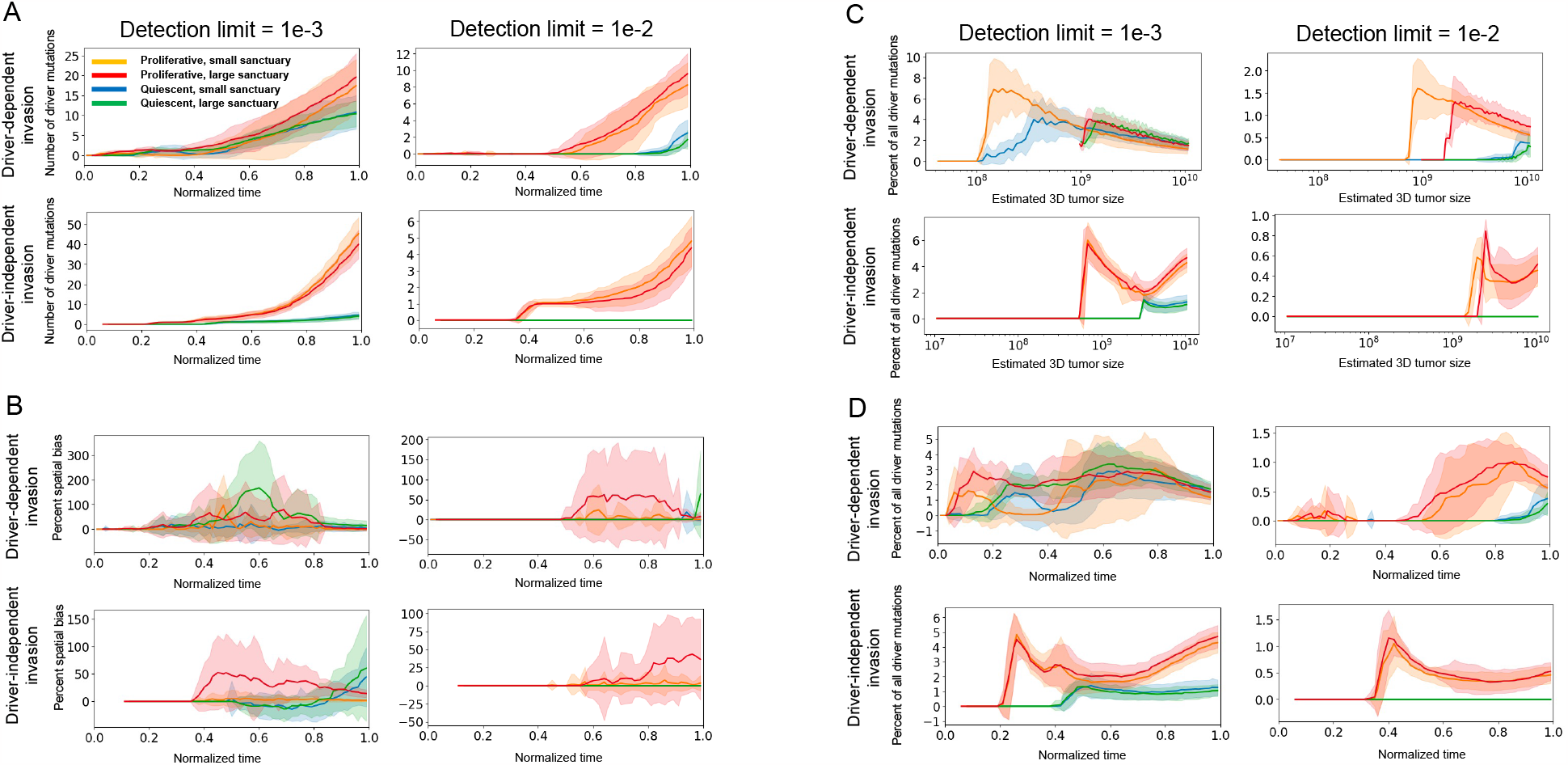
Number, percent spatial bias, and overall percentage of detectable drivers: **(A)** Plots of the number of detectable driver mutations starting from the point of relapse for minimum detection frequencies of 1e-3 and 1e-2, for proliferative and quiescent tumors, relapsing at ∼ 10^8^ and ∼ 10^9^ cells, over normalized timepoints. Mutations were detectable if the estimated VAF exceeded the detection limit. VAFs were estimated based on a tumor fraction of 1% for a 3 billion-cell tumor with death rate of 0.1 (see Methods). **(B)** Percent change in number of detectable drivers when the VAFs in (A) are compared to VAFs computed assuming the tumor sheds all clones at the same rate for the same detection limits, referred to as percent spatial bias (see Methods). **(C)** Overall percentage of detected driver mutations relative to population size. **(D)** Overall percentage of detected driver mutations relative to normalized timepoints.

**Figure S6.**
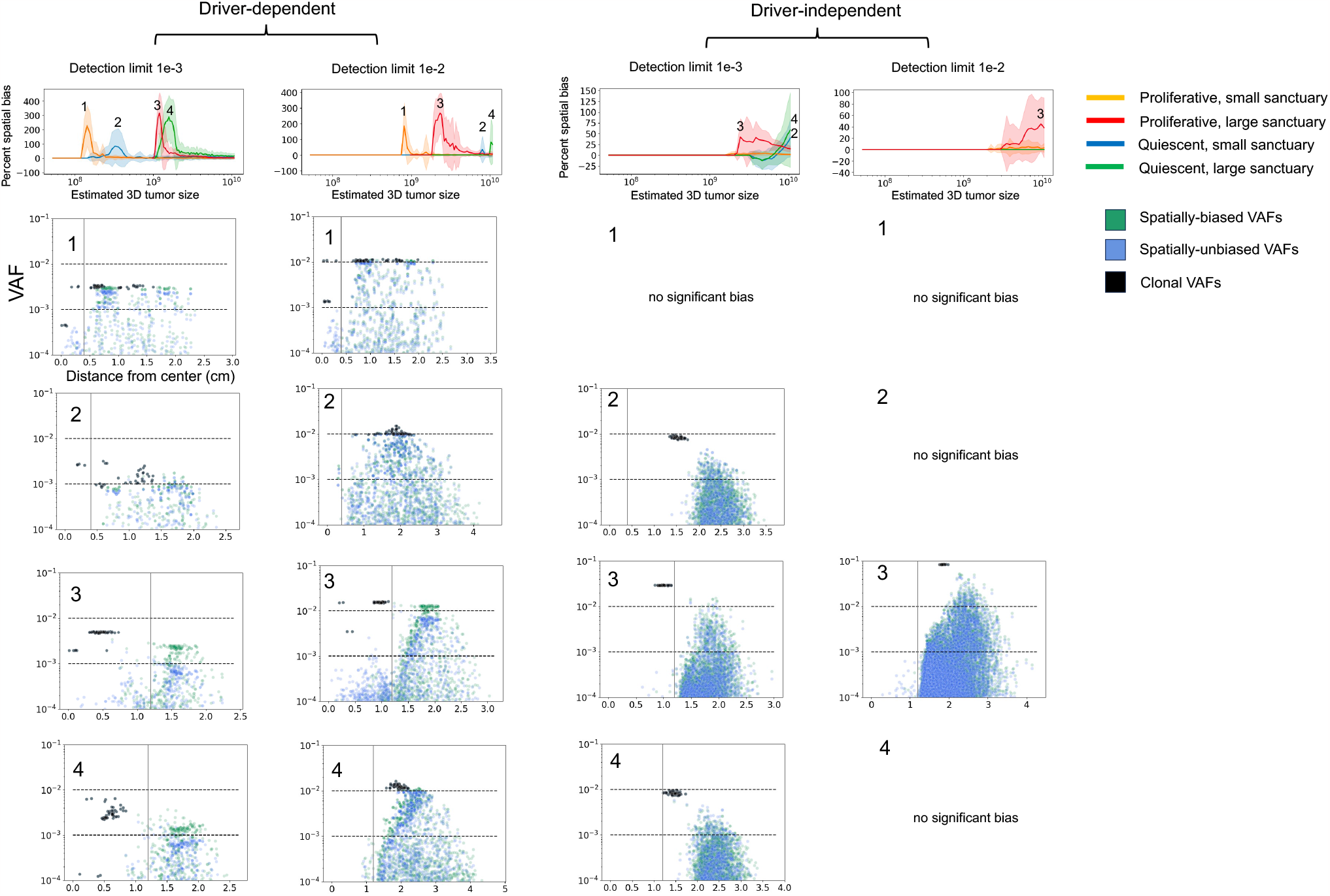
Spatial distribution of VAFs at points of maximal spatial bias for all scenarios: The top row of line plots are repeated from **Figure 4B**, showing the percent change in detected driver mutations for detection limits 1*e* − 3 and 1*e* − 2, under driver-dependent and independent invasion. Each scatterplot shows the distribution of VAFs corresponding to distance from the tumor center.

## Notes

### Competing Interest Statement

The authors have declared no competing interest.

